# Network modules linking expression and methylation in prefrontal cortex of schizophrenia

**DOI:** 10.1101/497891

**Authors:** Dongdong Lin, Jiayu Chen, Nora Perrone-Bizzozero, Jing Sui, Vince D. Calhoun, Jingyu Liu

**Affiliations:** The Mind Research Network and Lovelace Biomedical and Environmental Research Institute, Albuquerque, NM, 87106, USA; Dept. of Neurosciences, University of New Mexico, Albuquerque, NM, 87131, USA; Dept. of Electronic and Computer Engineering, University of New Mexico, Albuquerque, NM, 87131, USA; Dept. of Psychiatry, University of New Mexico, Albuquerque, NM, 87131, USA

**Keywords:** DNA methylation, gene expression, brain postmortem, and schizophrenia

## Abstract

Tremendous work has demonstrated the critical roles of genetics, epigenetics as well as their interplay in brain transcriptional regulations in the pathology of schizophrenia (SCZ). There is great success currently in the dissection of the genetic components underlying risk-conferring transcriptomic networks. However, the study of regulating effect of epigenetics, as a modulator of environmental factors, in the etiopathogenesis of SCZ still faces many challenges. In this work we investigated DNA methylation and gene expression from the dorsolateral prefrontal cortex (DLPFC) region of schizophrenia patients and healthy controls using weighted correlation network approaches. We identified and replicated two expression and two methylation modules significantly associated with SCZ. Among them, one pair of expression and methylation modules were significantly overlapped in the module genes which were enriched in astrocyte-associated functional pathways, and specifically expressed in astrocytes. Another two linked expression-methylation module pairs were involved aging process with module genes mostly related to oligodendrocyte development and myelination, and specially expressed in oligodendrocytes. Further examination of underlying quantitative trait loci (QTLs) showed significant enrichment in genetic risk of most psychiatric disorders for expression QTLs but not for methylation QTLs. These results support the coherence between methylation and gene expression in a network level, and suggest a combinatorial effect of genetics and epigenetics in regulating gene expression networks specific to glia cells in relation with SCZ and aging process.

## Introduction

Schizophrenia (SCZ) is known to be a highly heritable and developmental neuropsychiatric disorder, which has around 1% prevalence worldwide [1, 2]. In previous neuroimaging studies, SCZ has been characterized by brain-wide gray matter reduction [3], functional disconnectivity [4, 5] and myelination decrease [6, 7] along with significant impairments in cognition [8], and presence of positive and negative symptoms. These brain abnormalities are thought to be mediated by complex molecular processes involving neuronal cell development, differentiation and death, even during the early stage of brain development [9]. Despite substantial efforts on exploring the underlying molecular mechanisms through advanced genomics, epigenetics and transcriptomics techniques, there are still many challenges given the complex interplay among these factors for the etiology of SCZ.

Brain transcriptomics studies have demonstrated substantial transcriptional alterations in the frontal cortex, cerebellum or hippocampus of SCZ [10]. Keeping gene expression in balance is critical in maintaining neuronal function, as well as their synaptic interactions. Genetics is a major factor in regulating gene expression and functional pathways [11, 12]. Recent landmark work based on dorsolateral prefrontal cortex (DLPFC) postmortem tissue RNA sequencing has reported 20~50% of SCZ risk loci [13] showing strong cis-effect to their nearby genes’ expression, further elaborating the liability of these risk variants in etiology of SCZ [14, 15]. Co-expression network analyses of brain expression data have identified some modules highly associated with SCZ harboring the SCZ risk loci with strong cis-acting effects [15, 16]. However, some co-expression modules show discrepancy in relation with SCZ and the polygenic risk score for SCZ. Several SCZ-related modules fail to be significantly enriched in SCZ genetic susceptibility [14]. In addition, differentially expressed genes could have diverse patterns among brain regions [17], neural cells [18, 19] and neural developmental stages [20]. Collectively, these findings suggest the influence of complex interplay of genetics and environmental factors in determining the process of gene expression.

Epigenetics can mediate gene by environment effects in modifying how genes are structured and expressed. DNA methylation is an epigenetic modification widely studied in psychiatric disorders such as SCZ [21, 22]. It changes the genome’s response to transcriptional factors by attaching a methyl group in DNA sequence (mostly on cytosine site as CpG). The influence of DNA methylation in gene transcription can be stable and heritable across cell generations, and also reversible according to external condition changes [23]. The dynamics of DNA methylation plays a vital role in the pathogenesis of SCZ, especially in neuronal diversity, plasticity and neurogenesis [24, 25]. Besides environmental and developmental effects, DNA methylation can also be influenced by sequence variants (e.g., genotype variation or specific allele on a locus), which represents a methylation QTL (meQTL). Previous studies have demonstrated prevalent meQTL effects across genome, especially during the early neural development, and their relation to multiple psychiatric disorders including SCZ [26–28]. The study for common and tissue specific meQTL effects across brain and peripheral tissues shows that cross-tissue meQTLs are more likely to reside at regulatory elements (e.g., enhancer) of transcriptional genes and also highly enriched in eQTLs [29], suggesting the role of DNA methylation in mediating genetic effects on gene expression.

Although DNA methylation plays a very important role in the control of gene expression by modulating both genetics and environmental factors, there are currently limited number of studies directly investigating the relationship between DNA methylation and gene expression in the human brain. By leveraging public datasets of methylation and gene expression from brain DLPFC tissues, we performed a comprehensive network analysis with an aim to characterize the relationship between them. Given the co-regulation among genes in expression and the clustering of methylation in functional pathways, we sought to identify the relationship between functional transcriptional modules and epigenetic modules in the context of the human interactome. We applied network analyses to construct expression and methylation networks and prioritize them into multiple sub-network modules, followed by a systematic characterization of both types of modules in terms of their associations with SCZ and aging, neuronal cell specification, CpG-expression co-localization and their underlying genetic effects. Using these analyses, we were able to identify links between gene expression and methylation at the module level.

## Materials and Methods

### Gene expression data

The data were downloaded from dbGaP (Accession: phs000979.v1.p1). Briefly, brain postmortem tissues of DLPFC gray matter region (Brodmann area 46) from 546 subjects including SCZ and HCs were used for RNA extraction and gene expression assays. Further information about tissue dissection, clinical characterization, neuropathological screening, and toxicological analyses was described in [30]. SCZ patients were assessed for schizophrenia or schizoaffective disorder based on DSM-IV lifetime Axis I. HCs were included if they had no history of psychological or psychiatric problems and negative toxicology results.

Gene expression was assayed by an Illumina whole-genome HT-12-V4 expression chip covering 47323 probes. The transcript level of each probe was passed through quality control steps by the ‘limma’ R package [31], including correction for background level using negative control probes, quantile normalization using both negative and positive controls, and log_2_ transformation on the transcription intensities. Probes were removed if they were not significantly expressed (detection p-value > 0.01) in at least 90% subjects as applied in [32]. Subjects were excluded if they had age information missing or were less than 16 years old, or had poor RNA quality (RNA integrity number: RIN<6.5) or less than 10% of probes significantly expressed (quality detection p-value < 0.01). After these quality control steps, we had 20,370 probes from 419 subjects (253 HCs and 250 SCZ) left for analyses. Batch effects were corrected for each probe using a parametric Bayes framework implemented in the ‘combat’ function [33] in an R package ‘SVA’[34]. Confounding factors including pH, PMI (post-mortem interval) and RIN were further regressed out by a linear model prior to main analyses.

### DNA methylation data

The data were from NCBI GEO database (GEO database: GSE74193). Briefly DNAs from DLPFC region (BA9/46) of 244 subjects (108 SCZ and 136 HCs) with age greater than 16 years old were assayed using Illumina Infinium Methylation450k assay, covering 485,512 CpG sites. More details about the criteria for patient and controls recruitment and diagnosis can be seen in [26].

DNA methylation data went through a series of quality control steps performed by R package ‘minfi’ [35] as applied in [26, 36]. After applying quantile-based normalization to both methylated and unmethylated signals on each site, we calculated the beta values for subsequent analysis. CpGs were removed if they 1) coincided with SNPs or at single base extension [37], 2) located in non-specific probes [38], 3) contained more than 1% missing values (methylation values with detection p>0.05 were treated as missing values), or 4) were located on sex chromosomes. The remaining missing beta values were further imputed by KNN method as used in [39]. 377,698 CpGs were kept after preprocessing. Further CpG removal was applied if CpGs standard deviation (sd) was less than the measurement error standard deviation (sd = 0.047) estimated by 78 test-retest samples, resulting 61853 CpGs for analyses. Batch effects were then corrected for each CpG using the same steps as with expression data. Neuronal cell type proportions and the top four principle components of negative control methylation as estimated by [26] were regressed out for subsequent analysis.

### Network analysis

After preprocessing of both expression and methylation data, we constructed co-expression and co-methylation modules separately by using software WGCNA [40]. In brief, for each dataset the adjacency matrix was calculated by a power of 6 of the correlation matrix among nodes (i.e., expression probes and CpG) to have scale-free topology larger than 0.85, from which topology overlap matrix (TOM) was derived to measure connection similarity among nodes—the overlap between any two nodes in terms of the extent they were connected to the same other nodes in the network. Through the TOM matrix, an unsigned co-methylation/expression network was constructed and densely interconnected CpGs or expression probes were clustered into modules. Module eigengenes (ME), the first principal component of methylation or expression matrix in a module, were computed and tested for the association with SZ diagnosis, controlling for age, race and sex. Within a module, the correlation of each expression probe or CpG with ME was computed as the measure of their module membership (MM) [41], indicating how close a CpG or probe relates to the module. Each CpG or probe’s association with SZ diagnosis was also computed as group significance (GS), from which the correlation between MM and GS on each CpG or probe in the module was tested. The top CpGs or probes with both high MM and GS values were utilized to represent the module for demonstration.

### Overlap test between expression and methylation modules

After prioritizing probes or CpGs into different expression or methylation modules respectively, we assigned them with ensemble IDs based on genome assembly GRCh38 using R package ‘BioMart’ [42]. We only annotated the methylation sites to a gene and ensemble ID if it was located within the gene or nearby the gene transcription start site (TSS) (distance < 5k bps). For each pair of expression and methylation modules, we tested the significance of ensemble ID overlap by both two-sided Fisher’s 2×2 table exact test and permutation test. In Fisher’s exact test, given an expression module, we compared the odd of shared ensemble IDs with a methylation module to the odd of shared ensemble IDs with the other methylation modules. To account for the cases that some methylation sites were not annotated to any genes and the number of methylation sites varied by modules, we also applied a permutation method to reduce these influences in the test. For a specific expression module, the proportion of overlapped ensemble ID from a methylation module was taken as an observation. Then we generated null distribution by randomly sampling 10^5^ sets of methylation sites with the same module size from the whole methylation set. For each randomly selected methylation set, the proportion of ensemble ID included in the expression module was calculated. The percentage of sampled methylation sets having higher proportion than the observed proportion was taken as empirical p-value, denoted by P_perm. The significance levels from both methods were used in determining the significance of overlap between expression and methylation modules.

### Replication analyses

We used independent expression and methylation data sets to validate the findings from discovery data from three aspects: module’s association with phenotypes, module’s preservation, and probe-level associations of expression and methylation from linked modules.

*Expression replication dataset 1*(*ExpRep1*; GSE36192): postmortem samples from frontal cortex of 455 neurologically normal Caucasian subjects were assayed by HumanHT-12_v3 Expression BeadChips (48803 probes). More details about expression profiling were described in [43, 44]. After applying the same quality control and preprocessing (batch correction) as in discovery data, there were 343 subjects (age > 16) with 18,196 probes left for replication.

*Expression replication dataset2*(*ExpRep2*; GSE21138): postmortem tissues from DLPFC (BA 46) of 30 SCZ patients and 29 age- and sex-matched HCs [45] assayed by Affymetrix Human Genome U133 Plus 2.0 Array, covering 54675 probes, followed by normalization using dChip statistical model (DNA-Chip Analyzer), and log2 transformation as applied in [45] and batch correction by ‘SVA’[34]. Probes with present calls in at least one sample were left, resulting in 30,061 probes from 58 (30 SCZ and 28 HCs) subjects for analysis. Probes were assigned by gene symbols and matched with those from discovery expression data at gene level for replication.

*Methylation replication dataset1*(*MethyRep1; GSE61380*): 33 post-mortem brain DLPFC (BA9) samples (18 SCZ and 15 HCs) were obtained from Douglas Bell-Canada brain bank with more information about participant recruitment and diagnosis criteria introduced in [46]. DNA methylation was profiled using the Illumina Infinium HumanMethylation450K BeadChip and went through the same normalization, quality control, and imputation process as applied in discovery data to have 368,998 methylation sites remained for validation. Batch correction and five neural cell type proportions estimation were also applied using the same algorithms as used in discovery data analysis.

*Methylation replication dataset2* (*MethyRep2; GSE36194*): corresponding to expression replication dataset 1, methylation data from the same tissue of same subjects were profiled using with the Illumina Infinium HumanMethylation27K beadchip. The methylation data were preprocessed and quality controlled through the same steps as in discovery data analysis, resulting 26424 CpGs from 330 subjects for validation analysis.

#### Replication for modular’s disease and age relatedness

For those modules with ME vectors significantly associated with diagnosis, we validated the associations in the independent datasets. For expression modules, we used *ExpRep2* to select the probes from the same genes as the probes in the module and calculated the first PC of expression data for testing their associations with disease and age. A similar validation was applied for each significant methylation modules in *MethyRep1.* Replication significance level was set as 0.05.

#### Replication for module preservation

Three different measures were applied to validate if the properties of selected modules were preserved across different datasets. First, a local clustering coefficient is a normalized average number of all triangle connections associated with each node. The generalized clustering coefficient (GCC) by [47] for weighted network, accounting for weight information in each edge, was used to reflect the presence of the module structure and mean prevalence of connectivity around the nodes. A permutation test was applied to test the significance of GCC in replicated datasets. In each of 10^5^ runs, we randomly selected features (expression probes or methylation sites) from the entire set and calculated GCC to build the null distribution. P-value can be derived by the proportion of runs with larger GCC than the observed GCC. Second, a composite preservation statistic was previously proposed to combine the measures of module density preservation (i.e. to test if the connections in the module remain in an independent dataset) and module connectivity preservation (to test if the connectivity pattern is similar to that in the independent dataset) and tested in [48]. A permutation was performed to estimate the mean and variance of each measure (i.e., module density and module connectivity) under the null hypothesis that there was no preservation of module in the measures, followed by the standardization of each observed measure to the mean and variance under null distribution, denoted by Z_density and Z_connectivity, respectively. The composite preservation statistic, denoted by Z_summary, was defined as the average of Z_density and Z_connectivity. Z_summary < 2 indicates no preservation, 2<Z_summary<10 indicates weak to moderate evidence of preservation, while Z_summary>10 indicates strong evidence [48]. The p-value of the composite statistic was the combination of p values of module density and module connectivity.

#### Probe-level expression-methylation relation

To test directly regulation relation between expression and methylation at probe level for the linked modules, we leveraged the data from *ExpRep1* and *MethyRep2* that were collected from the same samples, and tested pair-wise associations of expression probes and methylation sites if they are located within 500k bps, using linear regression controlling for covariates age, sex and PMI.

### Neural cell type and functional enrichment

To understand the cell specificity of each expression and methylation modules, we tested their enrichment in the genes expressed specifically in five cell types (neuron, astrocytes, microglia, endothelia cells, and oligodendrocytes using the pSI R package [49] with Fisher’s exact test threshold as 0.05 and false discovery rate (FDR) as 0.05. The purified cell expression data from GSE73721 were used for the test [50]. The overlap of our modules with those cell-specific modules identified in a recent transcriptional study for psychiatric disorders [51] was also evaluated by Fisher’s exact test. In addition, methylation data of both neuron and glia cells from postmortem prefrontal tissues of 29 control subjects (GSE41826 [52]) were used to extract two sets of CpGs showing significant hyper-methylation in neuron (denoted by “neuron-up’) and glia cells (denoted by “glia-up”), respectively (p<1×10^−10^). These two sets of CpGs were then used to test the CpG overlap with identified methylation modules. To understand the biological function of each module, we performed a gene ontology (GO) and KEGG enrichment analysis on the genes in the module using the web tool Webgestalt (http://www.webgestalt.org/option.php).

### Psychiatric disorders GWAS risk loci enrichment

Public meQTL database (GSE74193) and eQTL databases (GTeX: https://gtexportal.org/home/; BrainSeq: http://eqtl.brainseq.org/phase1/devel) from brain DLPFC tissue were applied to access the genetic effects underlying expression and methylation networks. To test if the QTLs from target modules were significantly enriched in genetic risk loci of psychiatric disorders compared to the QTLs from the other modules, we adopted both Fisher’s exact test and permutation test as applied in [29]. Specifically, we firstly ran linkage disequilibrium (LD) pruning supervised by GWAS risk loci on the whole QTL set with r^2^>0.7 using LD structure from 1000 genome project EUR group as reference. After pruning, the proportion of QTLs from the target module showing GWAS risk was taken as the observation. Then we generated null distribution by randomly sampling 10^5^ sets of QTLs from the whole pruned QTL set with the same number of QTLs and similar minor allele frequency distribution as those from the target module. Empirical p-value was computed by the percentage of sampled QTL sets having higher proportion than the observed proportion, denoted by P_perm. Fisher’s exact test was used to test the odds ratio of risk loci in target module QTLs and the QTLs from the other modules, denoted by P_Fisher.

## Results

### Co-expression modules and their associations with SZ

We clustered expression probes into 22 modules using WGCNA software with the dendrogram plot as shown in Fig.1A. The eigengenes of each module were tested for association with multiple traits as shown in Fig.1B. Two modules colored by magenta (including 276 probes) and yellow (including 1098 probes) were identified to have significant associations with SCZ diagnosis (yellow: t-stat = 4.1, p = 2×10^-3^; magenta: t-stat = 3.55, p =9.7×10^-3^) after controlling for covariates and correction for multiple tests by false positive rate (FDR). For each SCZ related module, we tested the correlation between module membership and SCZ group significance for all probes within the module, as shown in Fig.S1 A-B (Supplementary file 1). It can be seen that the correlation is significant in both modules with values of 0.53 and 0.28, respectively, showing that the probes with higher disease association tend to be more important in representing the module.

**Figure.1.**
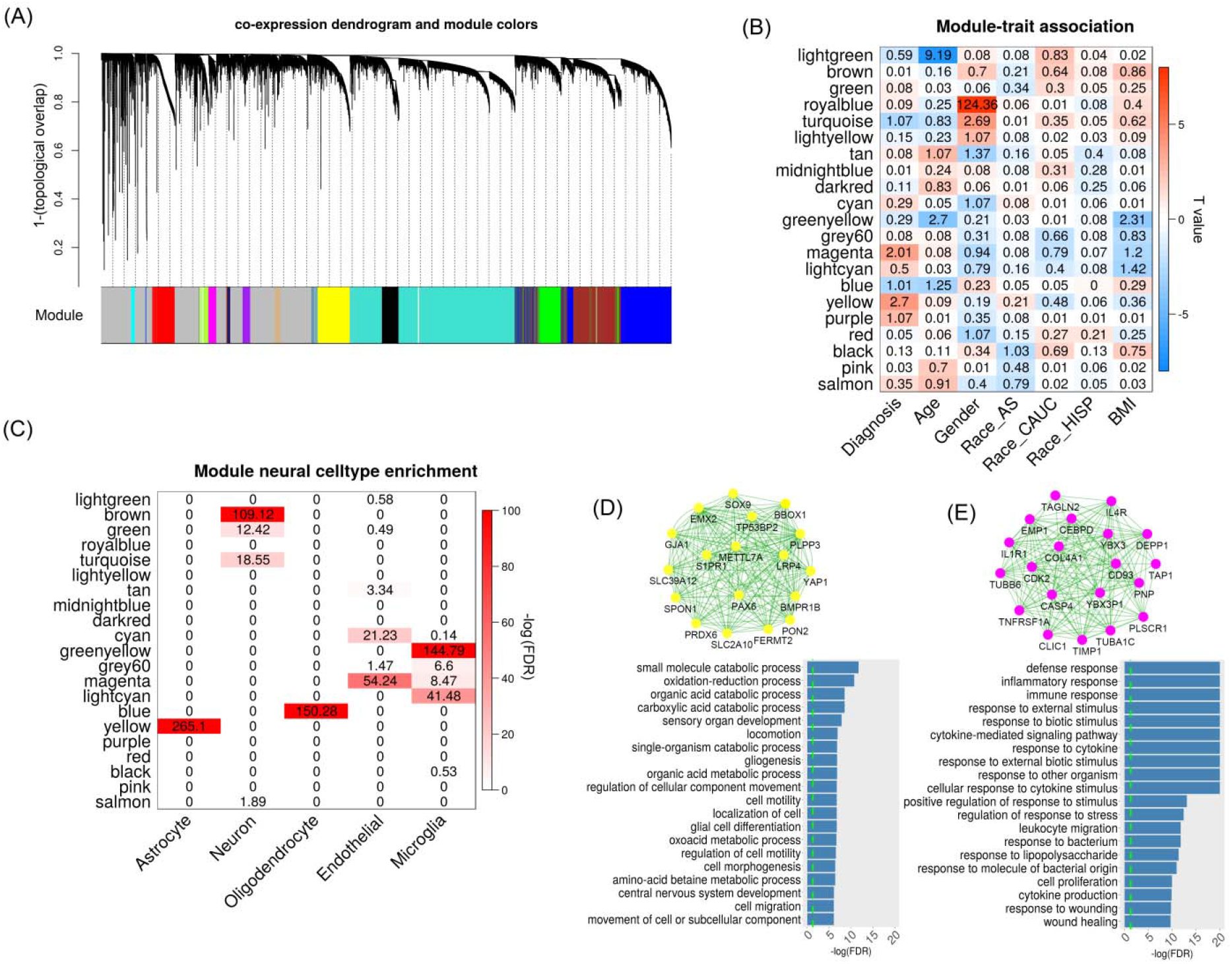
Co-expression analysis on brain DLPFC expression data. (A) shows the dendrogram clustering the correlated probes into several modules with different colors; (B) lists the relationship between module eigengenes and traits. The numbers indicate log_10_(FDR) and colors indicate the T-values. (C) shows the enrfichment of module genes in seven reported modules which are specific to neural cells (i.e., neuron, endothelial, astrocyte, microglia and oligodendrocyte). (D) and (E) plot the network for the top 20 hub genes and their functional enrichment in gene ontology for magenta and yellow modules, respectively.

Independent validations were applied to verify both SCZ-related modules in two datasets (*ExpRep1* and *ExpRep2*). In *ExpRep1* dataset, 90.8% of expression probes from the yellow module and 88.8% probes from the magenta module were identified for replication. By permutation tests, we found that both modules were well preserved with significantly larger GCC than random selection of probes (p<1×10^−4^, Fig.S3 A-B) and higher Z_summary statistics (yellow module: Z = 38; magenta module: Z = 18, Fig.S1 C). As recommended in [48], if Z_summary >10, there is strong evidence that the module is preserved. In *ExpRep2* dataset, 72.6% genes from the yellow module and 73.3% genes from the magenta module were identified. The expressions of those probes from the matched genes were used to calculate the first eigenvector for each module as module eigengene, followed by a test for schizophrenia association. We replicated significant disease associations of both modules with the same direction as identified in the discovery dataset (yellow module: t-stat = 2.54, p = 0.014; magenta module: t-stat =2.68, p = 9.6×10^-3^). In addition, both modules consistently showed significantly larger GCC in *ExpRep2* data (p<1×10^−4^, Fig.S3 C-D).

Several co-expression modules were significantly enriched in different neural cell types. For example, yellow expression module contains the genes mostly expressed in astrocytes while blue module is more specifically expressed in oligodendrocytes as shown in Fig. 1C. Gene ontology enrichment analysis identified significant enrichment of genes from the yellow module in the pathways mainly involved the cell growth, migration, differentiation, movement, and organ development and morphogenesis as shown in Fig.1D (FDR<0.05). Interestingly, they were also highly enriched in some neurodevelopment pathways such as gliogenesis, glial and astrocyte cell differentiation and central nervous system development. From Fig.1E, it can be seen that genes in the magenta module were mostly involved in the inflammatory response to external stimulus (e.g., cytokine and lipopolysaccharide), and were more specifically expressed in endothelial and microglia cells. Besides these modules, several others were also found to have significant enrichment in the cell-specific co-expression modules reported in a previous study [51], as shown in Fig.S4.

### Methylation modules and their associations with SZ

By a similar network analysis, we identified 13 methylation modules and tested the associations of these module eigengenes with multiple traits as shown in Fig.2A. There were three modules colored by yellow (641 CpGs, t-stat = −7.22, p =6.6×10^−11^), green (475 CpGs, t-stat = 3.78, p =7.5×10^−4^) and turquoise (15983 CpGs, t-stat = −2.87, p = 0.01) showing significant relationship with schizophrenia diagnosis (FDR<0.05). In each of three modules, CpGs tended to have more significant SCZ vs. HC group differences if they demonstrated higher membership within the module (correlations were 0.91, 0.61 and 0.43, respectively, Fig.S2 A-C). Fig.2C-D show the network structure of top 20 genes with high module membership in yellow and green modules. The genes from yellow module were significantly enriched in the functional pathways for developmental growth, axonogenesis, neurogenesis and neuron generations, while the genes from green module were related to cell projection, as shown in Fig.2C. Although genes in the turquoise module were also enriched in some neural development pathways such as forebrain development and central nervous system neuron differentiation, it had a large module size with less focus on specific functions and therefore was not further analyzed in this study. By matching the methylation nearby genes (<5kbp) with those from cell specifically expressed genes, we found strong enrichment of blue and red methylation modules in oligodentrocytes as shown in Fig.2B. Based on CpG patterns specific to neuron andglia cells, we identified that CpGs from blue and red methylation modules were more likely hypo-methylated in glia cells compared to neurons in human brain.

**Figure 2.**
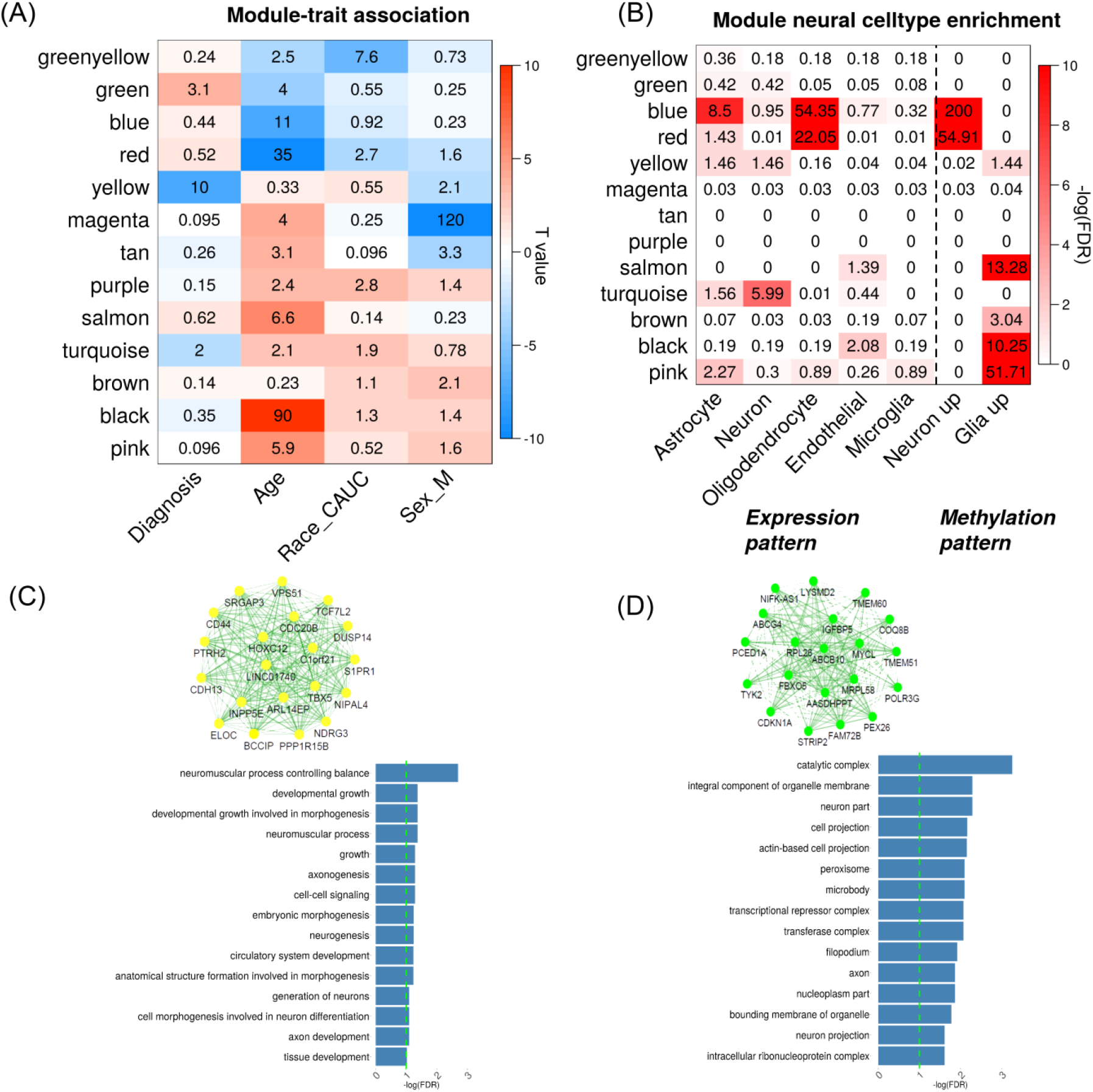
Network analysis on brain DLPFC methylation data. (A) shows the associations between module eigengenes and traits. The numbers indicate log_10_(FDR) and colors indicate the T-values. (B) lists the enrichment tests of each methylation module in different cell-specific modules; and (C-D) plot the network of top 20 representative genes and their functional enrichment in gene ontology for yellow and green module respectively.

To validate the above results in *MethyRep1* data, we matched 95.48% of CpGs in yellow module and 97.47% of CpGs in green module. We did not find any significant group difference in the first eigengene of yellow module (P = 0.28) but we found that the second eigengene showed group differences (t = −2.42, P = 0.02). The disease association for green module was significantly replicated with t = 2.98, P = 7.1×10^-3^. In addition, both modules were well preserved in the independent dataset with significantly higher GCC (P< 1×10^−4^, Fig.S3 E-F) and Z_summary statistics (yellow: Z = 13; green: Z = 21), as shown in Fig.S2 (D).

### Overlap between expression and methylation modules

After annotating methylation sites and expression probes to their target gene ensemble IDs, we tested the overlap among the expression and methylation modules based on both Fisher’s test (i.e. FDR_Fisher) and permutation test (i.e. FDR_perm). As shown in Fig.S5 (Supplementary file 1), we identified three sets of significantly overlapped modules which can be grouped to two categories: SCZ-related and age-related overlapping sets.

The SCZ-related overlapping set included the yellow expression module and yellow methylation module (odds ration OR = 1.75, FDR_Fisher = 0.04, FDR_perm = 0.009). Both modules were significantly associated with SCZ disease as shown in Fig.1B and Fig.2A. There were 1109 expression probes (nearby 893 genes; Supplementary file 2) and 641 CpGs (nearby 591 genes; Supplementary file 3) from each module, respectively, with 5.2% probes located closely (<5kbp) to 9.5% CpGs, denoted by ‘overlap CpGs’ and ‘overlap probes’. As shown in Fig.S6 (Supplementary file 1), overlap probes and CpGs were mainly involved the genes enriched in GO terms of regulating nitrogen compound metabolic process, biosynthetic process and fatty acid oxidation (FDR<0.05) and some interesting KEGG pathways like Wnt signaling, Adipocytokine signaling and Glutamatergic synapse(FDR<0.15). By annotating each CpG to its nearest gene, we found that 17.9% (p = 1.3×10^-3^) of CpGs were located within 100kbp of TSS of genes from the expression module, while 41.5% (p = 2.2×10^−3^) were within 500kbp, as shown in the density plot Fig.3A. Compared to those CpGs not included in the module, the overlap CpGs and the module CpGs(i.e., non-overlapped CpGs in the module) were significantly enriched in CpG islands (CGI) north shore (odds ratio: OR = 1.38 and 1.72), south shore (OR = 1.5 and 1.36), TSS200 regions (OR = 1.17 and 1.24), and 5UTR (OR = 1.34 and 2.18) as shown in Fig.3B.

**Figure 3.**
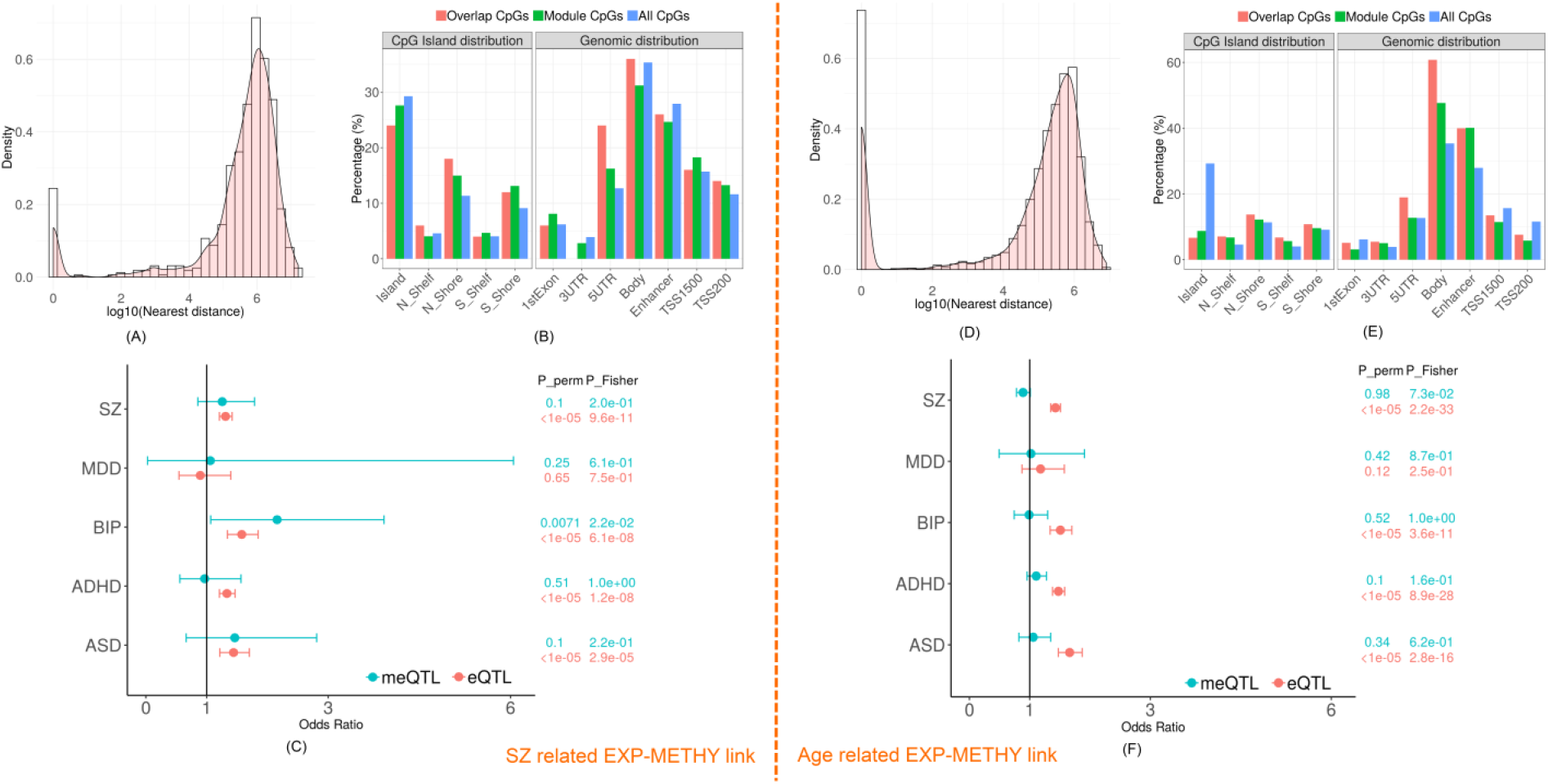
The characteristics of overlapping expression and methylation modules related to SCZ and age. (A) and (D) show the histogram plots of the distance between each methylation site and the nearest transcription start site of the linked modules. (B) and (E) plot the distribution of three sets of CpGs (overlap CpGs, module CpGs and all CpGs) across the genome. (C) and (F) list the enrichment tests of cis-meQTL and cis-eQTL in risk loci of five psychiatric disorders (SZ: schizophrenia, MDD: major depressive disorders, BIP: bipolar disorder, ADHD: attention deficit hyperactivity disorder, ASD: autistic disorder) from PGC study.

From previously reported eQTL and meQTL findings, we found that 32.8% of CpGs in the module (including 32% of overlap CpGs) were modified by cis-meQTLs, while 74% of expression probes (including 95.7% of overlap probes) were regulated by cis-eQTLs. And 12.6% of the cis-meQTLs are also cis-eQTLs for the expression module. In particular, 36.3% of cis-meQTLs modifying overlap CpGs, were also cis-eQTLs. In addition, 13 out of total 205 protein-coding genes in the methylation module were located in 108 SCZ risk regions [53] as listed in Table S1. The eQTLs also showed significant enrichment in risk loci of SCZ (OR = 1.31, p_perm <1×10^−5^; p_Fisher = 9.6×10^−11^) and three other psychiatric disorders (Autism spectrum disorder (ASD), Attention-deficit/hyperactivity disorder (ADHD) and dipolar disorder (BIP)) as shown in Fig.3C. None significant enrichment was found for meQTLs in any psychiatric disorders.

Two age-related overlapping sets were found. One was between the blue expression module and the blue methylation module (OR = 2.39, FDR_Fisher = 1×10^−39^, FDR_permutation <1×10^-5^), namely ‘blueExp-blueMethy’ link. The other was between the blue expression module and the red methylation module (OR = 5.79, FDR_Fisher =1×10^−30^, FDR_permutation <1×10^-5^), namely ‘blueExp-redMethy’ link. The blue expression module has a negative association with age as shown in Fig.1B (partial η^2^ = 0.025, p = 9.6×10^-3^) and the association was replicated in *ExpRep1* (89.6% probes included, p = 8.4×10^-3^) and *ExpRep2* (61.2% genes included, p = 4×10^-3^). The blue and red methylation modules were also negatively significantly associated with age as shown in Fig.2A (blue: partial η^2^ = 0.19, p = 1.2×10^-11^; red: partial η^2^ = 0.49, p = 2.5×10^-36^) and the age-methylation associations were also replicated (blue module: 96.6% CpGs included, p = 0.016; red module: 97% CpGs included, p = 2.3 ×10^-6^) in *MethyRep1.* GO term enrichment analysis showed that genes overlapped by blueExp-blueMethy modules were mainly enriched in functional pathways involving glial cell differentiation, gliogenesis, neurogenesis, axon ensheathment, and myelination (Fig.S7 (A)). Similar pathways were also significantly enriched by the overlapped genes from blueExp-redMethy linked modules, as shown in Fig.S7 (B).

In blueExp-blueMethy link, there are 2317 expression probes and 6308 CpGs with 20.9% CpGs located nearby 25.8% gene TSS (<5kbps). Fig.3D plots the distance distribution between CpGs and their nearest expression probes in the linked modules, showing that 36.4% CpGs were located within 100kbps of gene TSS and 65.5% CpGs were located within 500kbps. The overlap CpGs and module CpGs were mostly enriched in body of the gene and enhancer regions (OR = 1.77, p = 8.1×10^−100^ and OR = 1.85, p = 1.4×10^−107^, respectively) compared to all other CpGs, as shown in Fig.3E. In addition, 42.2% of CpGs from the module including 40.2% of overlap CpGs were significantly modified by cis-meQTLs, while 73.3% of expression probes including 94.9% of overlap probes were affected by cis-eQTLs. In particular, 46.2% of the cis-meQTLs are also cis-eQTLs for the expression module. Fig3.F shows the significance of eQTLs enriched in risk loci of SCZ, BIP, ADHD and ASD but none significant enrichment was found for meQTLs in any psychiatric disorders.

In the red methylation module (440 CpGs) which was also overlapped with blue expression module as shown in Fig.S8, 41.3% CpGs located closed to 5.4% of the expression gene TSS (<5kbps). 54.5% CpGs and 75.6% CpGs were located within 100kbps and 500kbps of TSS of genes from the expression module, respectively. CpGs from the red methylation module were significantly enriched in gene body (OR = 1.7, p = 1.5×10^-7^) and enhancer regions (OR = 2.8, p = 2.9×10^-27^) compared to all other CpGs. 54.5% of CpGs including 48.5% of overlap CpGs were modified by meQTLs, while 94.6% of overlap probes were affected by eQTLs. 35.7% of meQTLs were also cis-eQTLs in regulating the gene expression of the module.

Based on the *ExpRep1* and *MethyRep2* datasets assayed from the same subjects, we tested the associations between expression probes and their nearby (<500kbps) CpGs from each of the above overlapping sets. There were only few expression-methylation pairs (1.2~8.9%) matched due to lower resolution in *MethyRep2.* After controlling for covariates age, sex and PMI, 20%, 25% and 47.8% of matched pairs showed significant associations (FDR<0.05) in yellowExp-yellowMethy, blueExp-blueMethy and blueExp-redMethy links, respectively.

## Discussion

We used large transcriptomics and epigenetics data from human brain frontal cortex to explore the relationship between co-expression and methylation modules in terms of their overlap in genes, associations with SCZ and aging, as well as their shared underlying genetic risk. Our findings were replicated in multiple independent datasets on module’s associations with SCZ and age, and module’s preservation, and further verified by direct associations between individual CpGs and gene expression probes. We found two modules in each modality showing significant associations with SCZ, and three pairs of overlapping methylation and expression modules related to SCZ or aging.

### SCZ-related modules and their expression-methylation relationship

Two co-expression modules were significantly associated with SCZ with one (yellow module) including the genes more specifically expressed in astrocytes. This module is highly preserved across studies and also in line with the identified modules in previous RNA-sequencing studies of DLPFC tissue [15, 16, 51], showing consistent pattern with up-regulation of module expression in SCZ compared to HCs. In particular, the genes from the yellow module are also highly enriched in the module (CD4: astrocyte) reported in a recent study of transcription across five psychiatric disorders [51], which demonstrated significant changes of the module expression in SCZ, autism, and bipolar disorder. Astrocytes are known to be involved in synaptic metabolism and regulation of neurotransmitter release and reuptake (e.g., GABA and glutamate) and thereby critical for psychosis development[54]. Our pathway analysis found significant enrichment of the module genes in some SCZ-related KEGG metabolic pathways involving the metabolism on fatty acid [55], glycine [56], glutamate [57] and tryptophan [58], as shown in Fig.S9(A) (Supplementary file 1). Compared to the yellow module, the magenta module expression was also up-regulated in SCZ, but more specific to endothelial cells and microglia with genes mainly related with immune response. Previous studies have reported high involvement of immune system pathways in SCZ development via glia cells as well as their interactions with neurons in neurotransmitters perturbations [59]. The transcriptional changes of these immune system related genes suggest the complex co-expression patterns in downstream modulation of neuroinflammation in SCZ.

The yellow methylation module showed a significant association with SCZ. In particular, those methylation target genes included astrocyte highly-affinity glutamate transporters *SLC1A2* and *SLC1A3* [60], astrocyte-specific expressed marker gene *ALDH1L1* [61]. As shown in Fig.S9(B) and Table S2 (Supplementary file 1), they were also marginally enriched in GABAergic synapse (*GABRA1, GABRD, GNAO1, GNG7, PLCL1, PRKCA, CACNA1A*) and serotonergic synapse (*HTR1B, HTR2A, HTR3B, KCNJ3*), which are crucial neurotransmitter pathways in pathogenesis of SCZ [62–64]. Significant down-regulation of DNA methylation in most of CpGs near these genes (partial R^2^ = 1.8%~16.1%; Supplementary file 1, Table. S2) in SCZ, along with significant enrichment of module genes in growth development and neurogenesis, suggests the potential role of epigenetics in regulating glia cell function via these neurotransmitter signaling pathways. This is in agreement with previous findings of astrocyte epigenetic regulation in the pathophysiology of psychiatric disorders [54, 65, 66]. In addition, the genes nearby CpGs in the module were more likely to be mapped to PGC SCZ risk regions (p = 0.057) than the genes from the other methylation modules. In particular for SCZ high risk genes (Supplementary file 1,Table S1) including *RERE, MAD1L1, FURIN, GATAD2A,* they have been reported to show significant methylation level alterations in SCZ from previous DNA methylation genome-wide association studies [67–69].

The significant overlap between the yellow expression and the yellow methylation modules mainly involves genes with functions in some metabolic processes, biosynthetic process and fatty acid oxidation, which demonstrated the potential relationship with nervous system development in previous studies [61, 70]. In particular, CpGs nearby genes *SLC1A2, SLC1A3* and *PRKCA* from glutamate synapse showed hypo-methylation in SCZ which may lead to up-regulated expressions of these genes in astrocyte of SCZ, consistent with our results as shown in Table.S1 (Supplementary file 1). Significantly, a higher percentage of CpGs in the module were enriched in CGI shores which is known to be variable, compared the CpGs from the other modules. In addition, cis-eQTLs were identified for a large proportion of genes and enriched for SCZ and other three psychiatric disorders risk loci. These results suggest combinatorial effects of genetic and epigenetic components in regulating gene expression in astrocytes, dysregulation of which is associated with SCZ.

### Aging-related modules and their expression-methylation relationship

The module genes from blue expression module were highly enriched in genes expressed in oligodendrocytes, which are the CNS cells involved in myelination [71, 72]. GO enrichment test further validated that the functions of those specifically expressed module genes mainly involved oligodendrocytes differentiation, gliogenesis and development, ensheathment and myelination, which have been reported to be critical in maintenance of white matter integrity and structural connectivity in central nervous system [73, 74]. We found down-regulation of module expression, especially of the oligodendrocyte *OLIG2* gene (t = -2.56, p = 0.01) along with increase of age. *OLIG*2 can promote the formation of oligodendrocyte precursors and oligodendrocyte differentiation. Lower expression of *OLIG2* may cause deficits in oligodendrocyte production and differentiation and thus affect the formation of myelin, which is potentially related to the decreased ability of remyelination in the aging brain [75]. Although marginal association with SCZ was found in the first module eigengene, previous studies have shown the lower level of oligodendrocyte-associate gene expression in frontal cortex of SCZ along with myelination impairment and cognitive loss [6, 76]. Given that a number of studies have shown the dramatic loss of structural connectivity and white matter integrity across the brain regions in SCZ, it is promising to probe the role of oligodendrocytes-related expression module in the development of SCZ. In addition, the module harbored 13% SCZ risk genes and their cis-eQTLs were significantly enriched in most of psychiatric disorders risk loci, suggesting the potential pathology of genetic risk in SCZ through the alteration of module expression in oligodendrocytes.

Two methylation modules (blue and red) were significantly enriched in oligodendrocyte expressed genes and negatively associated with aging, pointing out the role of epigenetics in aging process by affecting oligodendrocyte development and differentiation. Previous work has reported the critical role of epigenetics in regulating oligodendrocyte gene expression and thereby leading to inefficient of remyelination during aging [77]. In our data DNA methylation of CpGs nearby myelin regulatory genes including *MYRF, MBP, MAG, MOG, CNP* [78], critical in oligodendrocyte development and myelination of axons, demonstrated significant negative associations with age (partial R^2^ = 2%~18%; Supplementary file 1, Table S3) in the blue module. CpGs nearby genes *MYRF*, *MBP* and *MAG* also showed significant decreases (partial R^2^ = 9%~39%; Supplementary file Table S3) along with the increase of age in the red methylation module. Additional pathway analyses showed that those myelin related genes together with many other genes in the modules were enriched for several functional pathways including central nervous development, neurogenesis, and glia cell differentiation, suggesting the potential of epigenetic regulations of these pathways in altering the oligodendrocyte differentiation and myelinations during aging process [79].

Functional pathways related to oligodendrocyte development and differentiation were shared by both expression and methylation modules, leading to significant overlap between the modules. There was a significant proportion of CpGs (20.9~41.3%) located nearby TSS of expression module genes (<5kbp), mostly enriched in gene body and enhancer regions, suggesting the regulation effect of those methylation module in expression modules [29]. In particular, by the association tests on the methylation and gene expression from the same subjects, we identified 25~47.8% of matched expression-methylation pairs mainly showing significant negative associations (Supplementary file 1, Table S4), including some oligodendrocyte highly associated genes *MYRF, MAG* and *SOX10.* Enrichment tests on neural specific methylation patterns showed that both methylation modules were enriched for hypo-methylation in glia cells compared to neurons, which might potentially contribute to their associated gene expression specificity in oligodendrocytes. These further evidences suggest epigenetic regulation on gene expression in oligodendrocytes to change myelination process. Genetics was also found to have large effects on both methylation and expression modules. A majority of module gene expressions were regulated by cis-eQTLs (74%~95%) and a relative high proportion of CpGs (40.2~54.5%) from both methylation modules were significantly regulated by cis-meQTLs. Especially, 35.7~46.2% meQTLs modifying overlap CpGs were also cis-eQTLs, indicating a pathway of some genetics in regulating oligodendrocyte gene expression through epigenetics. This genetic, epigenetic and expression interplay is in line with our previous knowledge of both gene and environmental factors involved in the process of demyelination during aging [80]. In addition, we found significant enrichment of cis-eQTLs in psychiatric disorders’ risk loci, which suggests shared genetics between risk for psychiatric disorders and regulating aging process which alter gene expression of oligodendrocytes, and partially reflect the similar decreases of myelination in both aging and SCZ patients.

To sum up, using a network analysis focused on DLPFC postmortem tissues, we identified and replicated two expression and two methylation modules significantly associated with SCZ with one pair showing significant overlap and enrichment in astrocyte-associated functional pathways. Additional overlapped expression-methylation module pairs were also replicated in relation to aging process with module genes mostly involved in oligodendrocyte development and myelination. Further examination of underlying QTLs showed the significant enrichment of eQTLs in most of psychiatric disorders’ risk loci, but not for meQTLs, suggesting combinatorial effects of genetics and epigenetics in regulating gene expression in glia cells of psychiatric disorders. This work as proof-of-concept demonstrates the existence of coherence between gene expression and DNA methylation at the network level, and provides support for the critical role of epigenetics in glial cell development and function during aging process and psychosis development.

### Limitations

The findings of this study should be interpreted in consideration of several limitations. Firstly, the analysis was performed on the datasets from different studies of different cohorts. Tests on the associations between methylation and expression from the same cohorts are still waiting for more efforts in psychiatric genetics research field. Although we tried to validate our findings in a study with both datasets, lower resolution of methylation array and missing of psychosis information limit our knowledge of the broad relationship between two features. Secondly, we have removed effects of some covariates (e.g., RNA quality, PMI, cell type and batch) prior network construction, but the modules may still be sensitive to the other factors (e.g., disease status and medication) and parameter settings in the method (e.g., block size and power index). Although we validated the preservation of identified modules in several independent datasets, more experiments are needed to evaluate the module robustness. Thirdly, genetic risk (eQTL and meQTL) were also discussed in the study, instead of computing eQTL from the data, we used the reports from public eQTL databases on RNAseq studies based on large sample size for eQTL detection. Although previous study has showed high overlap of RNAseq eQTLs with our microarray data [15], the results may still subject to the power and platform for eQTL detection.

